# Benchmarking predictions of MHC class I restricted T cell epitopes

**DOI:** 10.1101/694539

**Authors:** Sinu Paul, Nathan P. Croft, Anthony W. Purcell, David C. Tscharke, Alessandro Sette, Morten Nielsen, Bjoern Peters

## Abstract

T cell epitope candidates are commonly identified using computational prediction tools in order to enable applications such as vaccine design, cancer neoantigen identification, development of diagnostics and removal of unwanted immune responses against protein therapeutics. Most T cell epitope prediction tools are based on machine learning algorithms trained on MHC binding or naturally processed MHC ligand elution data. The ability of currently available tools to predict T cell epitopes has not been comprehensively evaluated. In this study, we used a recently published dataset that systematically defined T cell epitopes recognized in vaccinia virus (VACV) infected mice, considering both peptides predicted to bind MHC or experimentally eluted from infected cells, making this the most comprehensive dataset of T cell epitopes mapped in a complex pathogen. We evaluated the performance of all currently publicly available computational T cell epitope prediction tools to identify these major epitopes from all peptides encoded in the VACV proteome. We found that all methods were able to improve epitope identification above random, with the best performance achieved by neural network-based predictions trained on both MHC binding and MHC ligand elution data (NetMHCPan-4.0 and MHCFlurry). Impressively, these methods were able to capture more than half of the major epitopes in the top 0.04% (N = 277) of peptides in the VACV proteome (N = 767,788). These performance metrics provide guidance for immunologists as to which prediction methods to use. In addition, this benchmark was implemented in an open and easy to reproduce format, providing developers with a framework for future comparisons against new tools.

**Author summary:** Computational prediction tools are used to screen peptides to identify potential T cell epitope candidates. These tools, developed using machine learning methods, save time and resources in many immunological studies including vaccine discovery and cancer neoantigen identification. In addition to the already existing methods several epitope prediction tools are being developed these days but they lack a comprehensive and uniform evaluation to see which method performs best. In this study we did a comprehensive evaluation of publicly accessible MHC I restricted T cell epitope prediction tools using a recently published dataset of Vaccinia virus epitopes. We found that methods based on artificial neural network architecture and trained on both MHC binding and ligand elution data showed very high performance (NetMHCPan-4.0 and MHCFlurry). This benchmark analysis will help immunologists to choose the right prediction method for their desired work and will also serve as a framework for tool developers to evaluate new prediction methods.

## 1. Introduction

T cell epitope identification is important in many immunological applications including development of vaccines and diagnostics in infectious, allergic and autoimmune diseases, removal of unwanted immune responses against protein therapeutics and in cancer immunotherapy. Computational T cell epitope prediction tools can help to reduce the time and resources needed for epitope identification projects by narrowing down the peptide repertoire that needs to be experimentally tested. Most epitope prediction tools are developed using machine learning algorithms trained on two types of experimental data: binding affinities of peptides to specific MHC molecules generated using MHC binding assays, or sets of naturally processed MHC ligands found by eluting peptides from MHC molecules on the cell surface and identifying them by mass spectrometry. Since the first computational epitope prediction methods were introduced more than two decades ago [1–3], advancement in machine learning methods and increases in the availability of training data have improved the performance of these methods significantly in recent years, as has been demonstrated on benchmarks of MHC binding data [4, 5].

Given the wealth of epitope prediction methods available, it is necessary to keep comparing the performance of the different methods against each other, in order to allow users to rationally decide which methods to choose, and to allow developers to understand what changes can truly improve prediction performance. One issue with the past evaluations has been that, when new methods are developed and tested, they are commonly evaluated using the same kind of data on which they were trained, which can impact the performance results. For example, a method trained using MHC binding data will tend to show better performance when it is evaluated using MHC binding data and a method trained using MHC ligand elution data will tend to perform better when evaluated using MHC ligand data. The ultimate aim of the epitope prediction methods is to predict actual T cell epitopes i.e. peptides that are recognized by T cells in the host. Thus, we believe that the best way to compare prediction methods trained on different data is to evaluate their performance in identifying epitopes.

One problem when using T cell epitope identification as a way to benchmark prediction methods is that it is typically not known what a true negative is, as only a subset of epitope candidates is commonly tested for T cell recognition experimentally. Here, we took advantage of a recent study that comprehensively identified T cell responses in C57BL/6 mice infected with Vaccinia virus (VACV) [6]. This dataset is unique in that it covered all peptides previously shown to be presented by either H-2D^b^ or H-2K^b^ molecules expressed in these mice, which included epitopes identified following a large-scale screen of predicted peptide ligands [7], as well as all epitopes recognized in a comprehensive screen of a VACV protein expression library [8], and all peptides found to be naturally processed and presented by MHC ligand elution assays using mass spectrometry [6]. All these epitope candidates were rescreened in a consistent format, using eight separately infected mice, defining the major epitopes (categorized as those recognized in more than half of the animals), as well as negatives (never recognized in any animal), and for each epitope defining the magnitude of the T cell response.

We retrieved predictions from all publicly available computational algorithms prior to release of the dataset. We next evaluated each prediction algorithm based on its ability to pick the major epitopes from within the total peptides that can be derived from VACV, using different metrics such as AUC (area under the ROC curve), number of peptides needed to capture different fractions of the epitopes, number of epitopes captured in the top set of predicted peptides, and the magnitude of T cell response accounted for at different thresholds.

## 2. Materials & Methods

### 2.1 Selection of methods

As a first step, we compiled a list of all freely available CD8+ T cell epitope prediction methods by querying Google and Google Scholar. We identified 44 methods (S1 Table) that had executable algorithms freely available publicly (excluding those that required us to train a prediction model), and excluding commercial prediction tools that required us to obtain licenses. Out of these 44 methods, we selected those that had trained models available for the two mouse alleles for which we had benchmarking data (H-2D^b^ & H-2K^b^). Further, we contacted the authors of the selected methods and excluded the ones that the authors explicitly wanted to be excluded from the benchmarking for different reasons (mostly because the methods were not updated recently or new version of the methods were to be released soon). The final list included 15 methods that were selected to be included in the benchmarking: ARB [9], BIMAS [2], IEDB Consensus [7], MHCflurry [10], MHCLovac [11], NetMHC-4.0 [12], NetMHCpan-3.0 [13], NetMHCpan-4.0 [14], PAComplex [15], PREDEP [16], ProPred1 [17], Rankpep [18], SMM [19], SMMPMBEC [20], SYFPEITHI [3]. Out of the 15 methods, NetMHCpan-4.0 offered two different outputs, the first one being the predicted binding affinity of a peptide (referred as NetMHCpan-4.0-B), and the second the predicted probability of a peptide being a ligand in terms of a probability score (NetMHCpan-4.0-L). Both these outputs were evaluated separately. Similarly, MHCflurry could use two different models, first one trained with only binding data (MHCflurry-B) and second one incorporating data on peptides identified by mass-spectrometry (MHCflurry-L). Both these models were evaluated separately. Considering these as separate methods, a total of 17 methods were included in the benchmark, and are described in more detail in S1 Table. The methods differed widely in the peptide lengths that they could predict for each allele. For example, while MHCLovac could predict lengths 7-13 for both alleles, PAComplex could predict for only 8-mers of H-2K^b^ and none of the lengths in case of H-2D^b^. The methods also differed in the kind of prediction scores provided but ultimately they all represented a score that was intended to correlate with the probability of a peptide being an epitope in the context of the given MHC molecule. A complete list of the peptide lengths allowed for prediction per allele by each method and the kind of prediction scores they provide are given in S2 Table.

### 2.2 Dataset of VACV peptides

For the benchmark analysis, we used the peptide data set described in Croft et al., 2019 (S3 Table). This dataset represented a comprehensive set of peptides naturally processed and eluted from VACV-infected cells in addition to any previously identified epitopes. The total of 220 VACV peptides were tested for T cell immune responses in infected mice. Of these peptides, 172 were eluted from H-2D^b^ and K^b^ molecules from VACV-infected cells as described in detail in Croft et al., 2019. In brief, DC2.4 cells (derived from C57BL/6 mice [21] that expressed H-2^b^ MHC molecules were infected with VACV. The H-2D^b^ and K^b^ molecules were then individually isolated and the bound peptides eluted. The peptides were then analyzed by high resolution liquid chromatography-tandem mass spectrometry (LC-MS/MS). The remaining peptides in the set were not detected by LC-MS/MS and included 46 VACV-derived H-2^b^ restricted peptides/epitopes from the IEDB [22] and one entirely unpublished epitope and another that was mapped from a longer published sequence [23] identified by the Tscharke laboratory. Immune reactivity for each of these 220 peptides was then tested 8 times and the peptides that tested positive more than four times were classified as “major epitopes” and those tested positive four or fewer times were classified as “minor epitopes”. All peptides that were never positive were classified as “nonimmunogenic”. There were 83 peptides classified as “major” positives (S3 Table), ranging in lengths 7-13. In addition to the 220 peptides tested for immunogenicity, we generated all possible peptides of lengths 7-13 from the VACV reference proteome (https://www.uniprot.org/proteomes/UP000000344) (S4 File), which were also considered non-immunogenic, based on them not being found in elution assays on infected cells, and not being found positive in any of the many studies recorded in the IEDB. The entire dataset comprised 767,788 peptide/allele combinations.

### 2.3 Performance evaluations

The performance of the prediction methods was evaluated mainly by generating ROC curves (Receiver operating characteristic curve) and calculating the AUC_ROC_ (Area under the curve of ROC curve). The ROC curve shows the performance of a prediction model by plotting the True positive rate (TPR, fraction of true positives out of the all real positives) against the False positive rate (FPR, fraction of false positives out of the all real negatives) as the threshold of the predicted score is varied. AUC_ROC_ is the area under the ROC curve which summarizes the curve information and acts as a single value representing the performance of the classifier system. A predictor whose prediction is equivalent to random will have an AUC = 0.5 whereas a perfect predictor will have AUC = 1.0. That is, the closer the AUC is to 1.0, the better the prediction method. AUC values were first calculated on different sets of peptides grouped by length and allele separately. Secondly, overall AUCs were calculated by taking peptides of all lengths and both alleles together, which reflects the real life usability of having to decide which peptides to test. In this calculation, poor scores were assigned to peptides of lengths where predictions were not available for a given method. For example, in the case of SMM, lower numerical values of the prediction score indicate better epitope candidates, with scores ranging from 0 to 100. So a score of 999 was assigned to all peptides of lengths for which predictions were not available in SMM (lengths 7, 12 and 13 for both alleles). Similarly a score of −100 was assigned in case of SYFPEITHI (H-2D^b^: 7-8, 11-13; H-2K^b^: 7, 9-13) where a higher numerical value of predicted score indicates better epitope candidate and the scores ranging from −4 to 32.

### 2.4 Fully automated pipeline to generate benchmarking metrics

The Python scikit-learn package [24] was used for calculating the AUCs and Python matplotlib package [25] was used for plotting. A python script that can generate all results and plots along with the input file containing all peptides and their prediction scores from each method, immunogenicity category, T cell response scores, the “ProteinPilot confidence scores” representing the mass-spectrometry (MS) identification confidence level of the peptides and the number of times the peptides were identified by MS are provided in the GitLab repository (https://gitlab.com/iedb-tools/cd8-t-cell-epitope-prediction-benchmarking). The repository also contains the outputs from the script, i.e. the relevant results and plots. This will enable interested users to check our results and add their own prediction algorithms.

## 3. Results

### 3.1 Performance of the methods based on AUC_ROC_

As described in the method section, we identified 17 distinct prediction approaches that were freely accessible and could be applied to our dataset. Predictions from these methods were retrieved for all peptides of lengths 7-13 in the VACV proteome, which included the peptides tested for T cell response in Croft et al. (2019) [6]. The predictions were done using default parameters and the prediction outputs were used as provided by the tools without any modification or optimization. For tools provided by DTU server (NetMHCpan, NetMHC) and IEDB (Consensus, SMM, SMMPMBEC, ARB), where it provides raw score (for example predicted absolute binding affinity) and the percentile ranks (predicted relative binding affinity), the percentile ranks were used in the analysis. We considered the “major epitopes” (peptides that were tested positive in more than four out of the eight mice) as positives. To avoid ambiguity we excluded the “minor epitopes” (peptides that were tested positive in four or less mice out of the eight), and all other peptides were considered as negatives. This provides a binary classification of peptides into epitopes/non-epitopes. In order to evaluate the performance of each prediction approach, we generated ROC curves and calculated the AUC_ROC_ for all methods on a per allele (H-2D^b^, H-2K^b^) and per peptide length (7-13) basis, which are listed in Table 1. The per allele/length AUCs were then averaged to get an AUC value per each allele for each method and then the AUCs of both alleles were averaged to get a single AUC value per method. These average AUC values for each method are also provided in Table 1. The average AUCs varied from 0.793 to 0.983. NetMHCpan-4.0-B came top based on this analysis with an average AUC of 0.983. It was followed by NetMHCpan-3.0 (AUC = 0.982) and NetMHC-4.0 (AUC = 0.980). The lowest AUC was obtained for MHCLovac (0.793). When looking at the individual AUC values for each length, it was noticed that MHCLovac had very low performance for H-2K^b^ lengths 7 and 12 (AUC of 0.529 and 0.284 respectively) where there were only one positive each. Thus, these two low AUCs brought the average AUC down for MHCLovac, which is arguably irrelevant, as there are very few peptides positive for those lengths in the first place.

**Table 1.**
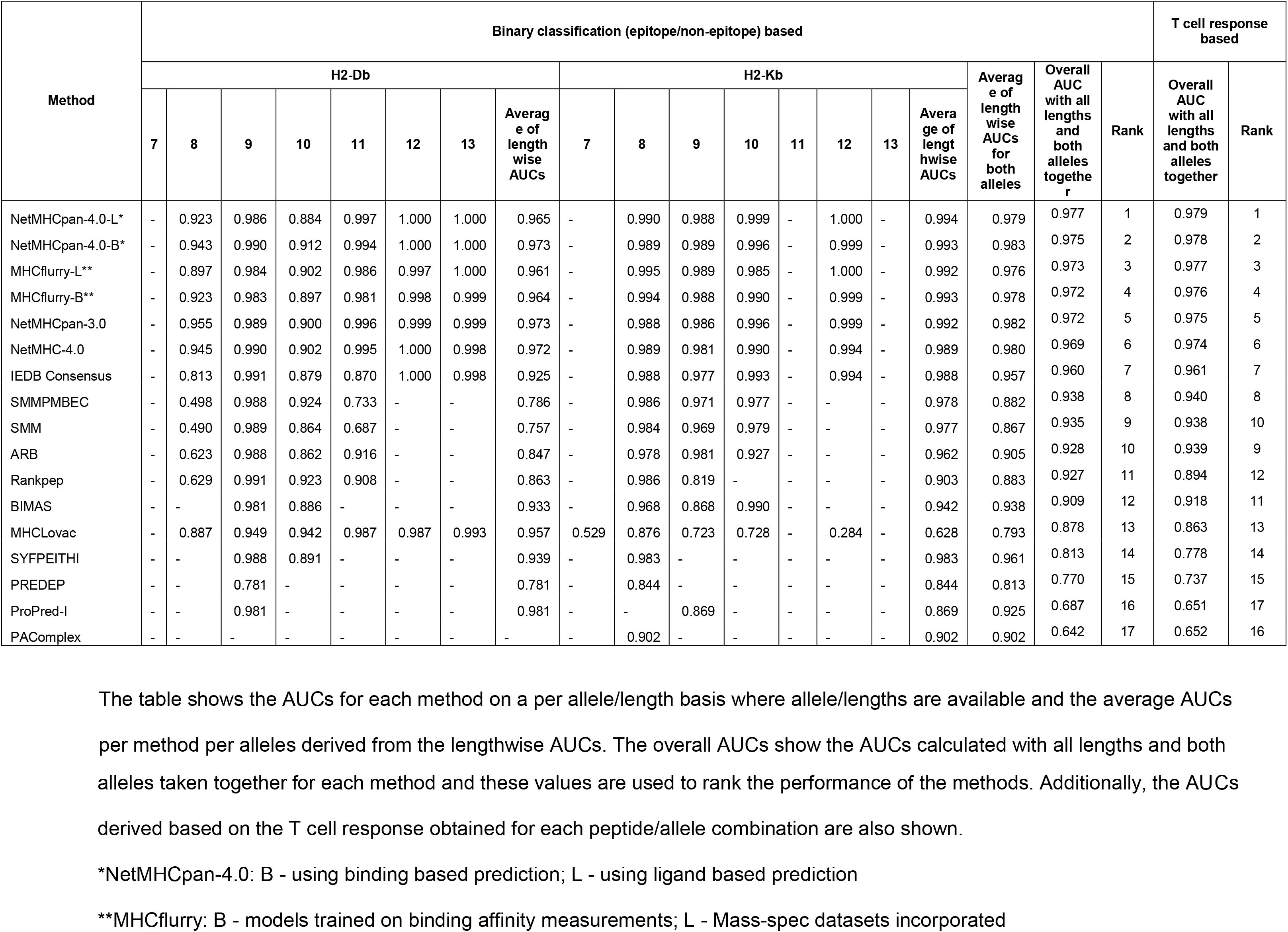
AUCs showing performance of each benchmarked method.

In practical applications, an experimental investigator uses predictions to choose which peptides to synthesize and test. The total number of peptides to be synthesized and tested is the limiting factor, and how many of the epitopes are covered is a measure of success, regardless of what the peptide length is or what allele they are restricted by. To reflect this, we estimated overall AUC values for each method by considering peptides of all lengths and both alleles together. If a given prediction method was unable to make predictions for a certain length (reflecting that the length is not considered likely to be an epitope), uniformly poor scores were assigned to those peptides. The overall AUCs ranged from 0.642 to 0.977. NetMHCpan-4.0-L ranked first with with AUC of 0.977 followed by NetMHCpan-4.0-B (0.975) and MHCflurry-L (0.973) (Table 1, Fig 1A). The ROC curves are shown in Fig 2. Fig 2A shows the ROC curves of all benchmarked methods for 100% FPR and Fig 2B shows the same up to 2% FPR to clearly distinguish the curves for each method in the initial part. Fig 2C and 2D show respectively the same for a set of top and historically important methods. It has to be noted that certain methods such as NetMHCpan-4.0 are implicitly adjusting prediction scores to account for the fact that certain peptide sizes are preferred when natural ligands are considered, as these methods were trained on such ligands. This means that prior approaches to adjust for the prevalence of different peptide lengths as was done for NetMHCPan 2.8 [26] are no longer necessary for such modern methods. It is likely that other methods, such as BIMAS or SMM that were trained on binding data only, could be improved when adjusting for lengths, but we wanted to test the performance of each method on an as-is basis.

**Fig 1.**
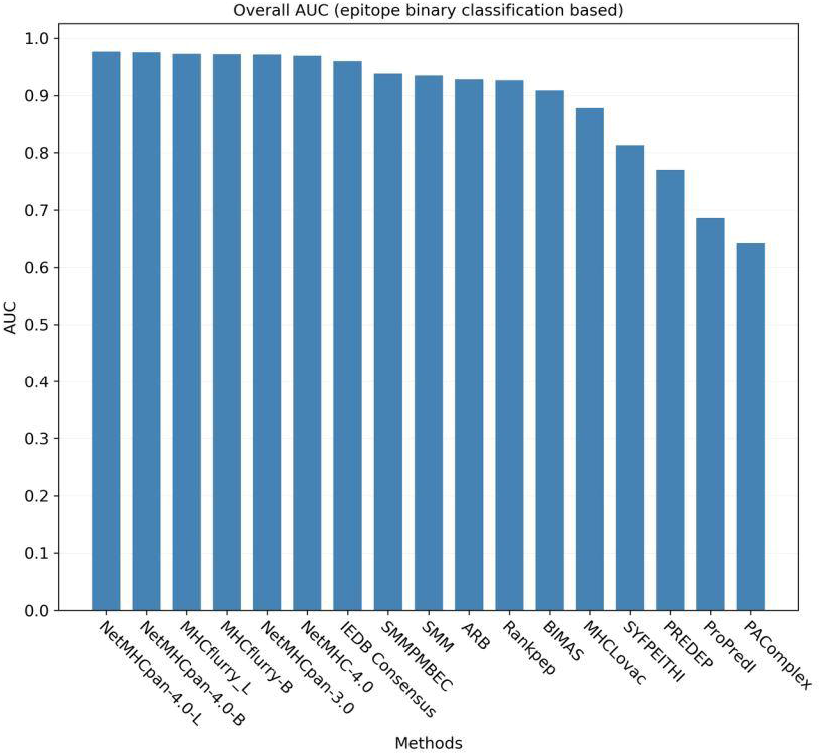
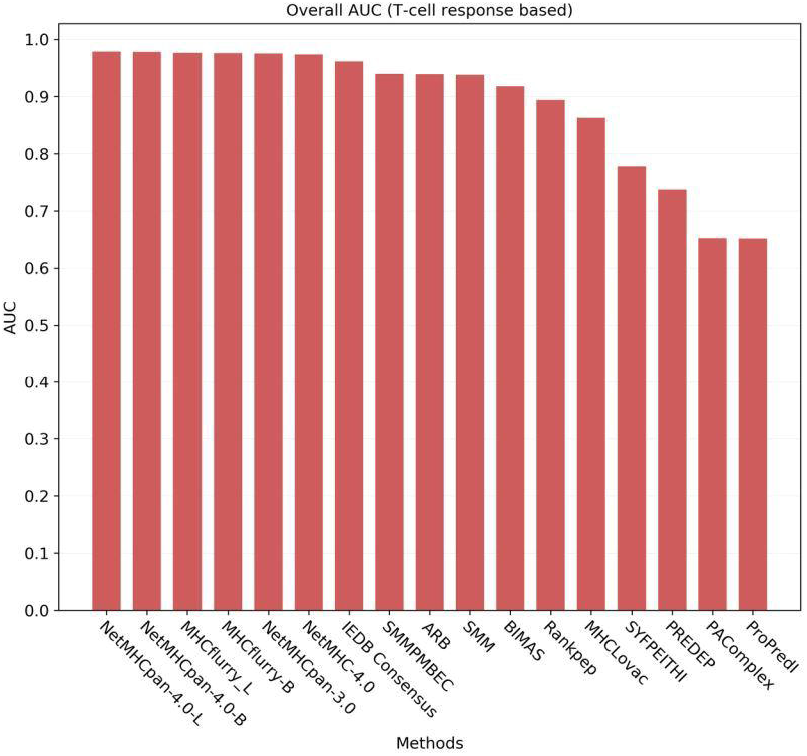
Bar charts showing the overall AUCs for each benchmarked method. Fig 1A. Bar chart showing the overall AUCs for each method with a binary classification (epitope/non-epitope) based analysis Fig 1B. Bar chart showing the overall AUCs for each method with a T cell response based analysis

**Fig 2.**
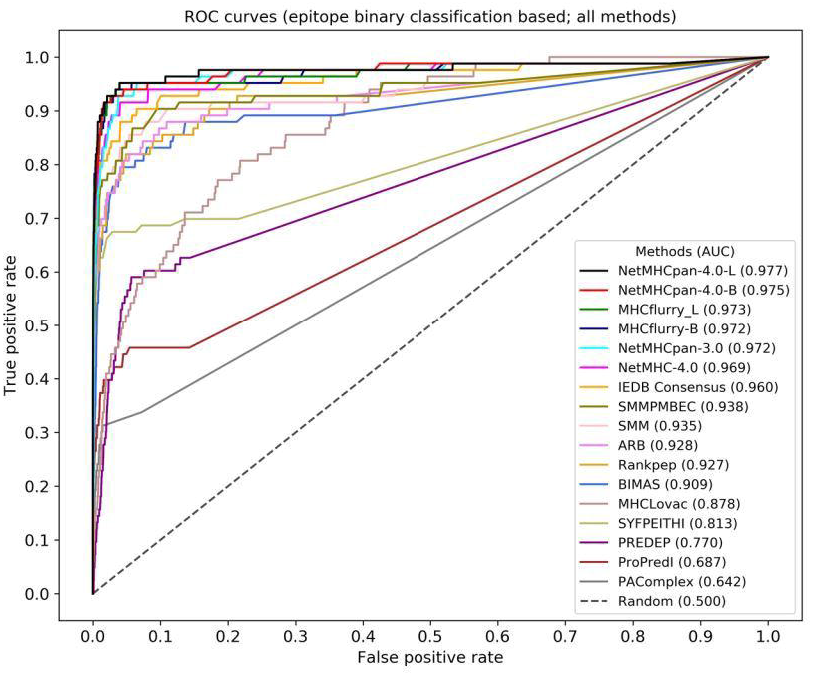
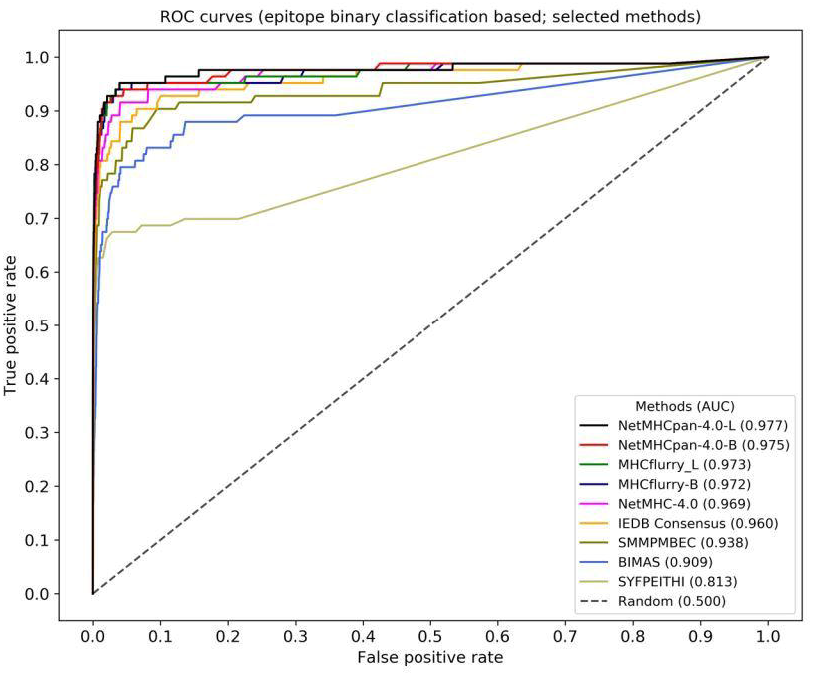
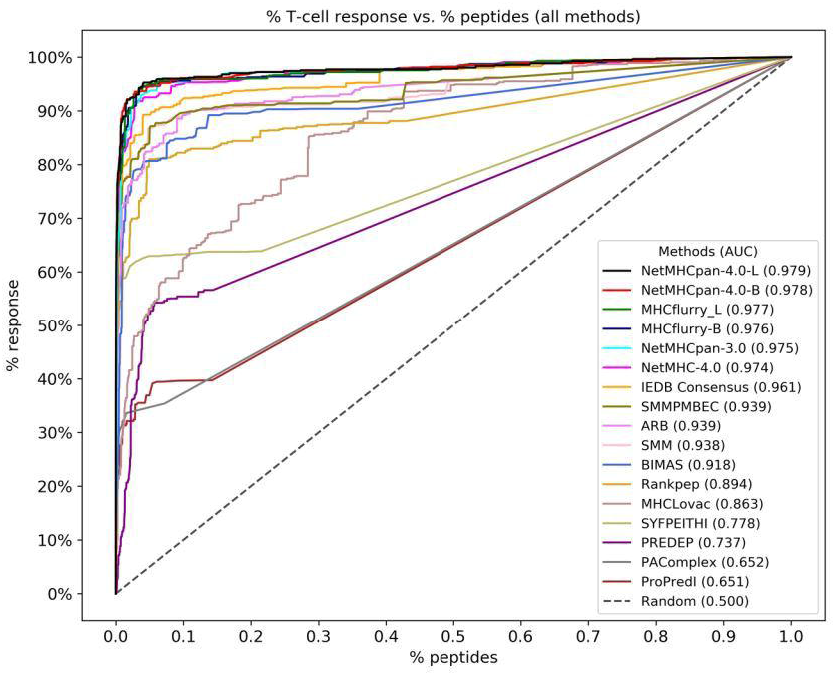
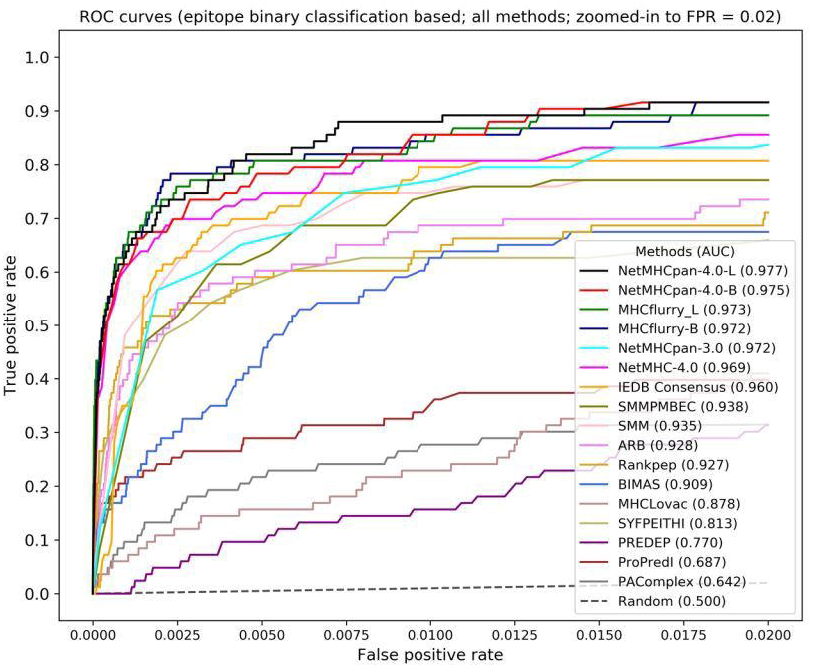
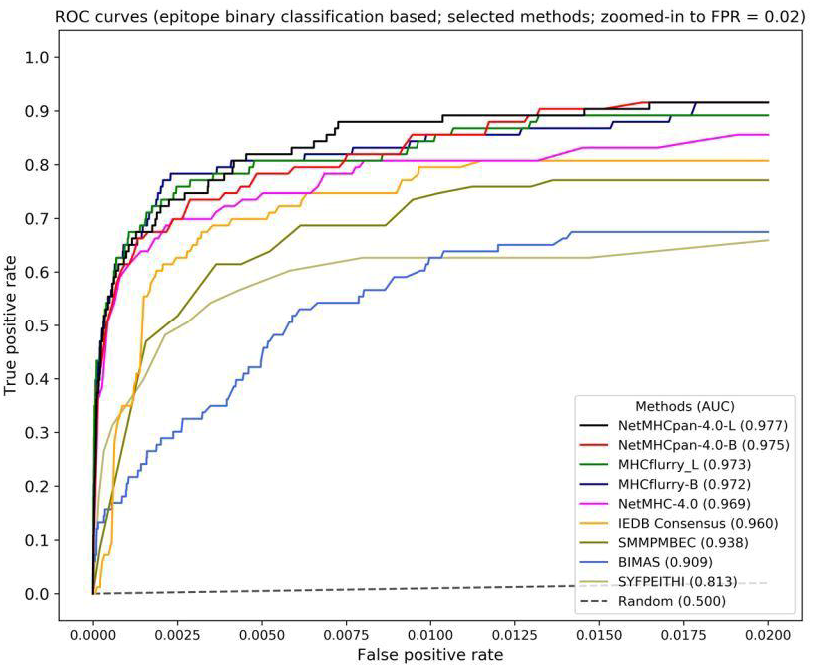
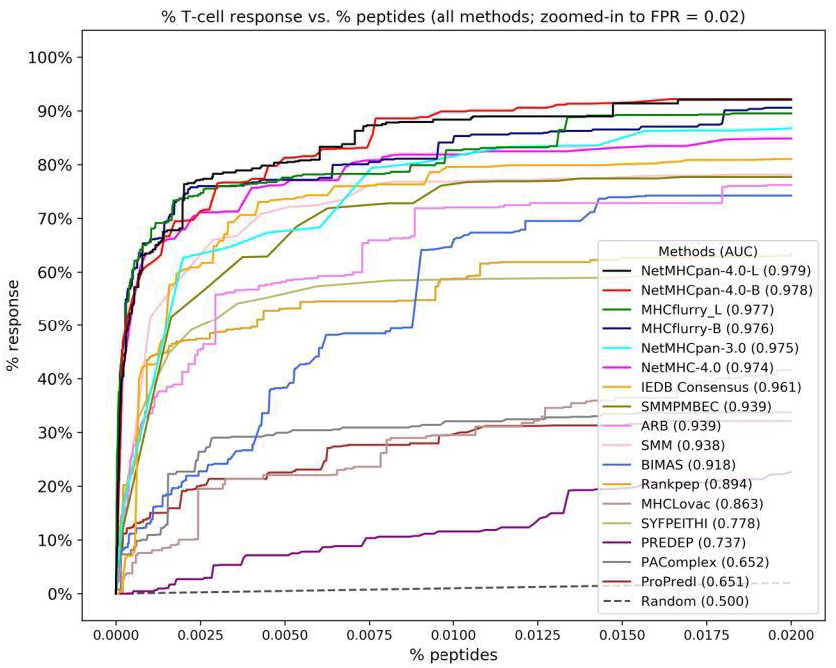
ROC curves showing the performance of the benchmarked methods. The curves are made by plotting true positive rate against the false positive rate in case of binary classification (epitopes/non-epitopes) based analysis and by plotting the % of T cell response against % of total peptides in case of T cell response based analysis. Fig 2A. ROC curve for all methods that were benchmarked. Fig 2B. ROC curve for all methods that were benchmarked with the curves zoomed in to FPR = 0.02 in order to be able to distinguish them more clearly in this region. Fig 2C. ROC curve showing the performance of a set of top and historically important methods. Fig 2D. ROC curve for selected methods with the curves zoomed in to FPR = 0.02. Fig 2E. Curve generated by plotting the % of T cell response against % of total peptides. Fig 2F. Curve generated by plotting the % of T cell response against % of total peptides. This plot shows the curves zoomed in to % of peptides = 0.02.

### 3.2 Alternative metrics to evaluate performance of the methods

In addition to the AUCs, we calculated metrics that are more intuitive for scientists less familiar with ROC curves, namely the number of peptides needed to capture 50%, 75% and 90% of the epitopes (which corresponds to comparing ROC curves at horizontal lines at 50%, 75% and 90% sensitivity). Since a total of 83 major epitopes were found in the dataset, we calculated how many predicted peptides were needed to capture 42 (= 50%) of them, after sorting based on the prediction score for each method. The results are shown in Table 2 and Fig 3A. The number of peptides required by the methods varied widely. NetMHCpan-4.0-L required only 0.036% (N = 277) peptides and MHCflurry-L needed only 0.037% (N = 285) peptides to capture 50% epitopes while ProPred1 needed 21% (160,644) and PAComplex needed 30% (230,132) peptides respectively to capture 50% epitopes. In a similar manner, we also calculated the number of peptides needed to capture 75% (N = 62) and 90% epitopes (N = 75). For 75% epitopes, MHCflurry-B was on top with 0.20% peptides (N = 1,542) whereas PAComplex needed 65% peptides (N = 498,917) (Table 2, Fig 3B). For 90% epitopes NetMHCpan-4.0-B needed only 1.33% (N = 10,224) peptides and NetMHCpan-4.0-L required only 1.47% (11,254) peptides while ProPred1 and PAComplex needed 84% (N = 646,291) and 86% (660,189) peptides respectively (Table 2, Fig 3C).

**Fig 3.**
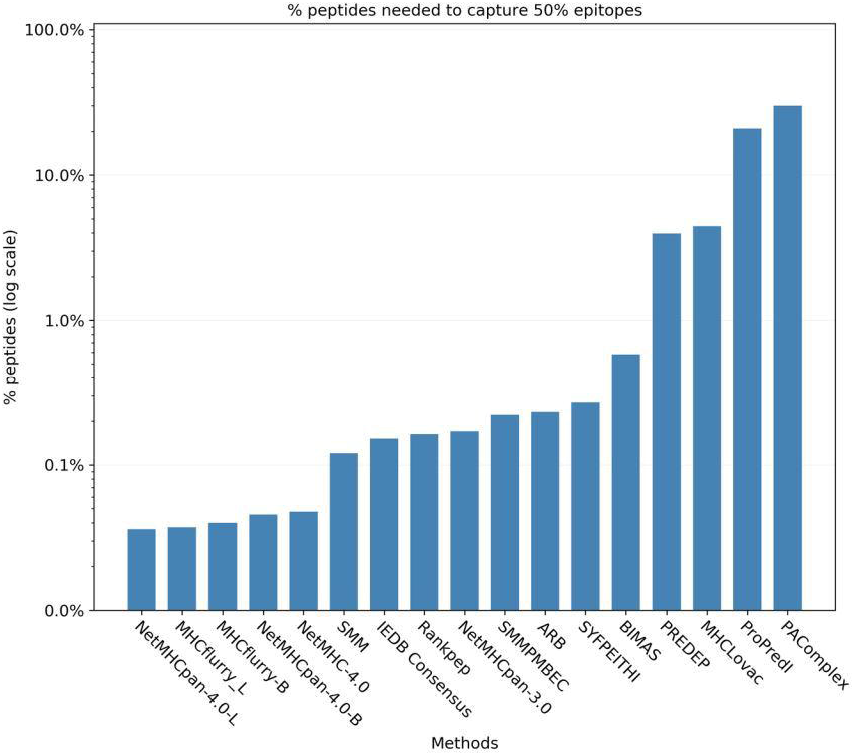
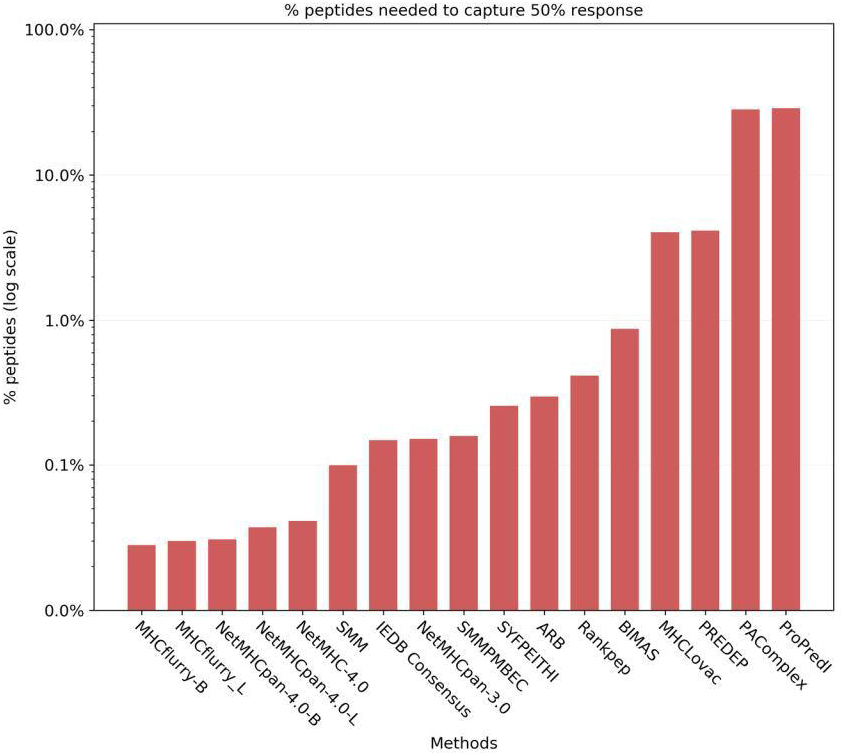
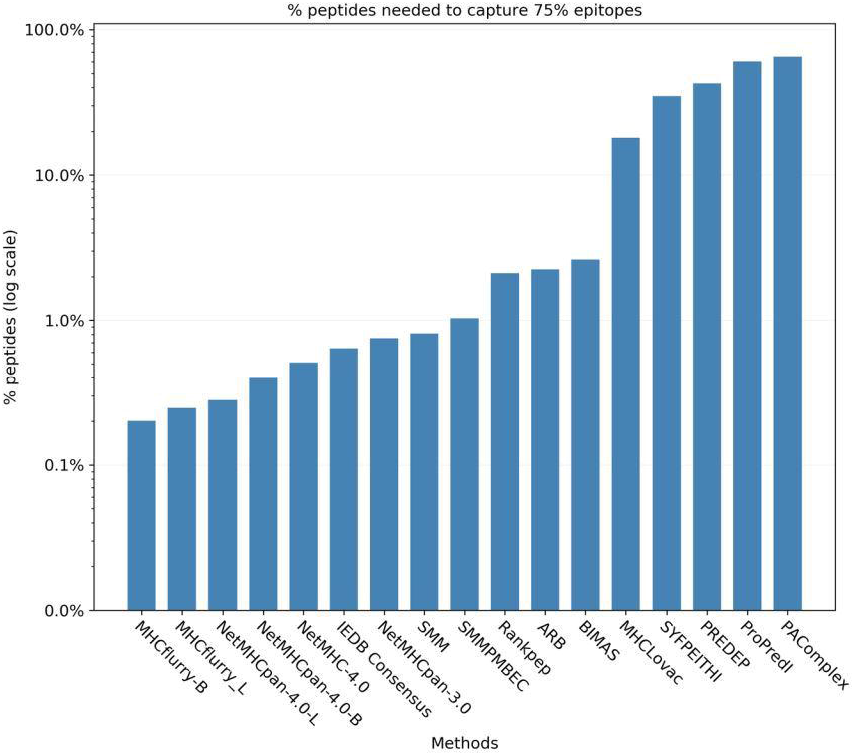
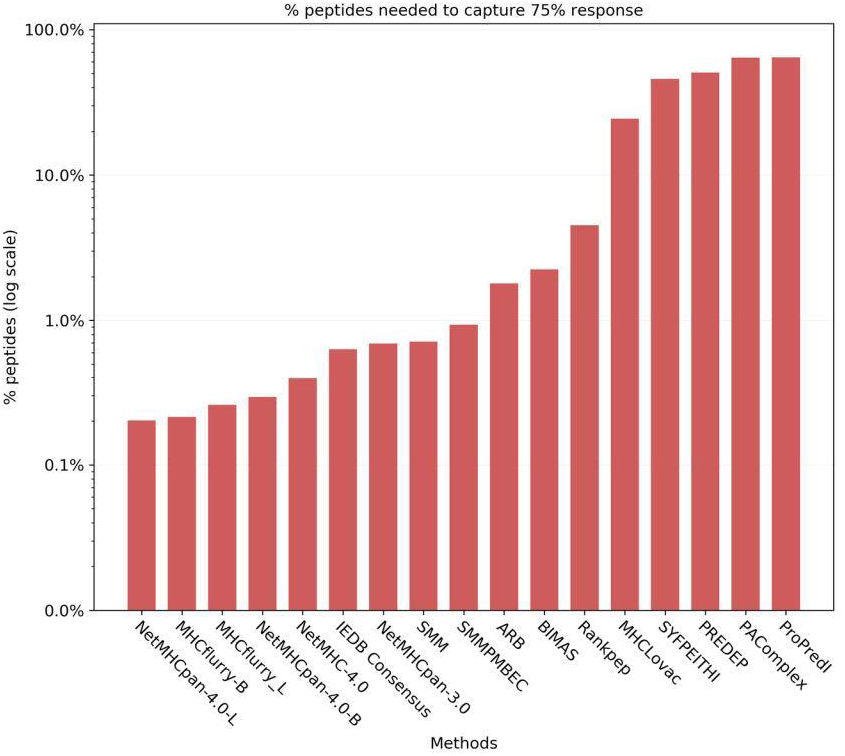
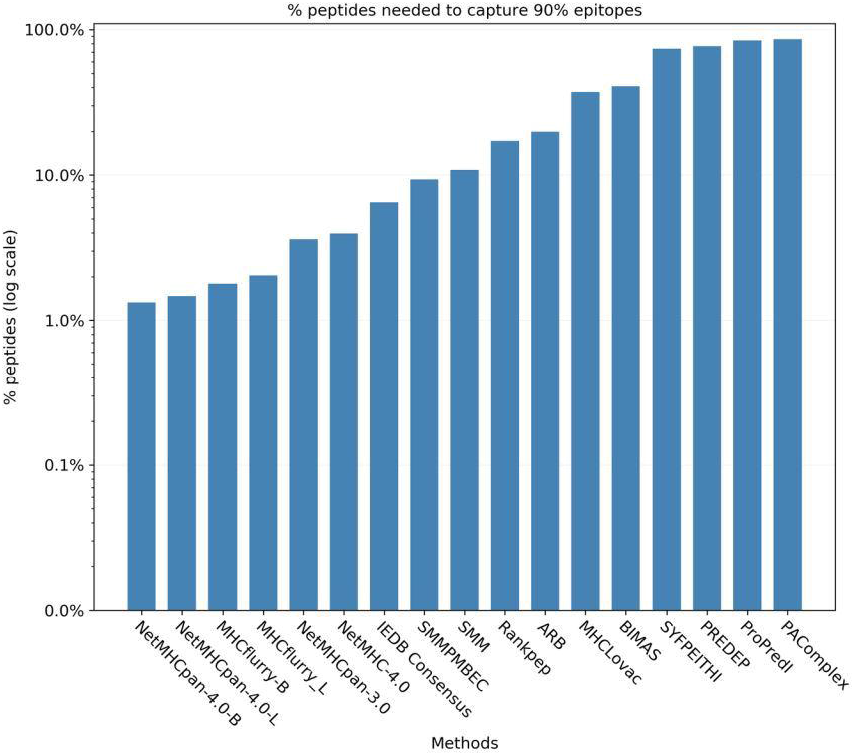
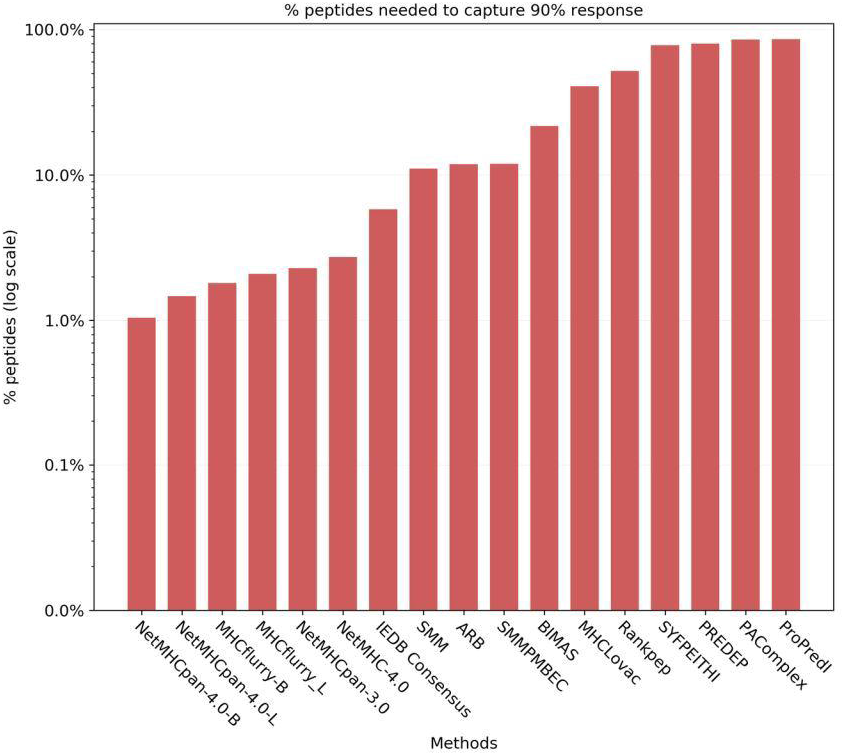
Number of peptides needed to capture 50%, 75% asnd 90% epitopes and T cell response. Fig 3A. Number of peptides needed to capture 50% epitopes. Fig 3B. Number of peptides needed to capture 75% epitopes. Fig 3C. Number of peptides needed to capture 90% epitopes. Fig 3D. Number of peptides needed to capture 50% T cell response. Fig 3E. Number of peptides needed to capture 75% T cell response. Fig 3F. Number of peptides needed to capture 90% T cell response.

**Table 2.** Number of peptides needed to capture 50%, 75% and 90% of epitopes and T cell response.

| Method | Peptides needed to capture 50% |  |  |  | Method | Peptides needed to capture 75% |  |  |  | Method | Peptides needed to capture 90% |  |  |  |
| --- | --- | --- | --- | --- | --- | --- | --- | --- | --- | --- | --- | --- | --- | --- |
|  | Epitopes |  | T cell response |  |  | Epitopes |  | T cell response |  |  | Epitopes |  | T cell response |  |
|  | Count | % | Count | % |  | Count | % | Count | % |  | Count | % | Count | % |
| NetMHCpan-4.0-L | 277 | 0.04% | 286 | 0.04% | MHCflurry-B | 1,542 | 0.20% | 1,639 | 0.21% | NetMHCpan-4.0-B | 10,224 | 1.33% | 8,030 | 1.05% |
| MHCflurry-L | 285 | 0.04% | 230 | 0.03% | MHCflurry-L | 1,896 | 0.25% | 1,991 | 0.26% | NetMHCpan-4.0-L | 11,254 | 1.47% | 11,309 | 1.47% |
| MHCflurry-B | 307 | 0.04% | 216 | 0.03% | NetMHCpan-4.0-L | 2,147 | 0.28% | 1,549 | 0.20% | MHCflurry-B | 13,719 | 1.79% | 13,842 | 1.80% |
| NetMHCpan-4.0-B | 349 | 0.05% | 236 | 0.03% | NetMHCpan-4.0-B | 3,058 | 0.40% | 2,250 | 0.29% | MHCflurry-L | 15,651 | 2.04% | 16,039 | 2.09% |
| NetMHC-4.0 | 365 | 0.05% | 317 | 0.04% | NetMHC-4.0 | 3,922 | 0.51% | 3,037 | 0.40% | NetMHCpan-3.0 | 27,731 | 3.61% | 17,533 | 2.28% |
| SMM | 924 | 0.12% | 761 | 0.10% | IEDB Consensus | 4,925 | 0.64% | 4,877 | 0.64% | NetMHC-4.0 | 30,472 | 3.97% | 20,984 | 2.73% |
| IEDB Consensus | 1,163 | 0.15% | 1,135 | 0.15% | NetMHCpan-3.0 | 5,764 | 0.75% | 5,341 | 0.70% | IEDB Consensus | 49,777 | 6.48% | 44,516 | 5.80% |
| Rankpep | 1,251 | 0.16% | 3,211 | 0.42% | SMM | 6,240 | 0.81% | 5,493 | 0.72% | SMMPMBEC | 71,593 | 9.33% | 91,619 | 11.93% |
| NetMHCpan-3.0 | 1,309 | 0.17% | 1,157 | 0.15% | SMMPMBEC | 7,939 | 1.03% | 7,174 | 0.93% | SMM | 83,425 | 10.87% | 84,821 | 11.05% |
| SMMPMBEC | 1,697 | 0.22% | 1,214 | 0.16% | Rankpep | 16,218 | 2.11% | 34,742 | 4.53% | Rankpep | 131,992 | 17.19% | 399,634 | 52.05% |
| ARB | 1,781 | 0.23% | 2,262 | 0.29% | ARB | 17,260 | 2.25% | 13,791 | 1.80% | ARB | 152,456 | 19.86% | 91,256 | 11.89% |
| SYFPEITHI | 2,070 | 0.27% | 1,955 | 0.25% | BIMAS | 20,156 | 2.63% | 17,264 | 2.25% | MHCLovac | 285,408 | 37.18% | 312,869 | 40.75% |
| BIMAS | 4,466 | 0.58% | 6,733 | 0.88% | MHCLovac | 138,245 | 18.01% | 187,337 | 24.40% | BIMAS | 313,329 | 40.81% | 166,819 | 21.73% |
| PREDEP | 30,363 | 3.96% | 31,820 | 4.14% | SYFPEITHI | 267,557 | 34.85% | 351,034 | 45.72% | SYFPEITHI | 567,644 | 73.94% | 601,086 | 78.29% |
| MHCLovac | 34,218 | 4.46% | 30,981 | 4.04% | PREDEP | 327,655 | 42.68% | 388,964 | 50.66% | PREDEP | 591,684 | 77.07% | 616,259 | 80.26% |
| ProPred1 | 160,644 | 20.93% | 221,775 | 28.89% | ProPred1 | 464,173 | 60.46% | 494,782 | 64.44% | ProPred1 | 646,291 | 84.19% | 658,585 | 85.78% |
| PAComplex | 230,132 | 29.98% | 216,523 | 28.19% | PAComplex | 498,917 | 64.99% | 492,155 | 64.10% | PAComplex | 660,189 | 86.00% | 657,535 | 85.64% |

The table shows the number of peptides needed to capture 50%, 75% and 90% of epitopes and T cell response. The lower the number of peptides needed to capture the respective amount of epitopes or T cell response, the better the performance of the prediction method.

Similar to above, another metric we calculated was the number of epitopes captured in the top 172 peptides predicted by each method. This corresponds to the number of peptides identified by mass-spectrometry of naturally eluted ligands. The results are provided in Table 3 and Fig 4A. The number of epitopes captured by these top peptides also varied widely for the methods. The MHCflurry methods performed the best, capturing 43% (N = 36) of the epitopes and NetMHCpan-4.0 methods captured 40% (N = 33) epitopes while PREDEP could not capture any epitope in the top 172 peptides.

**Fig 4.**
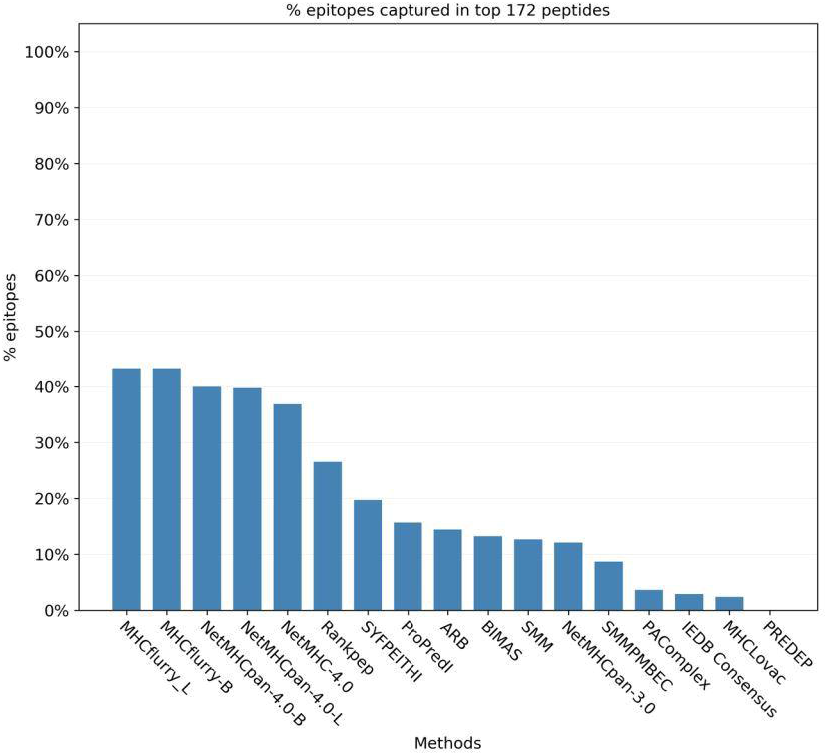
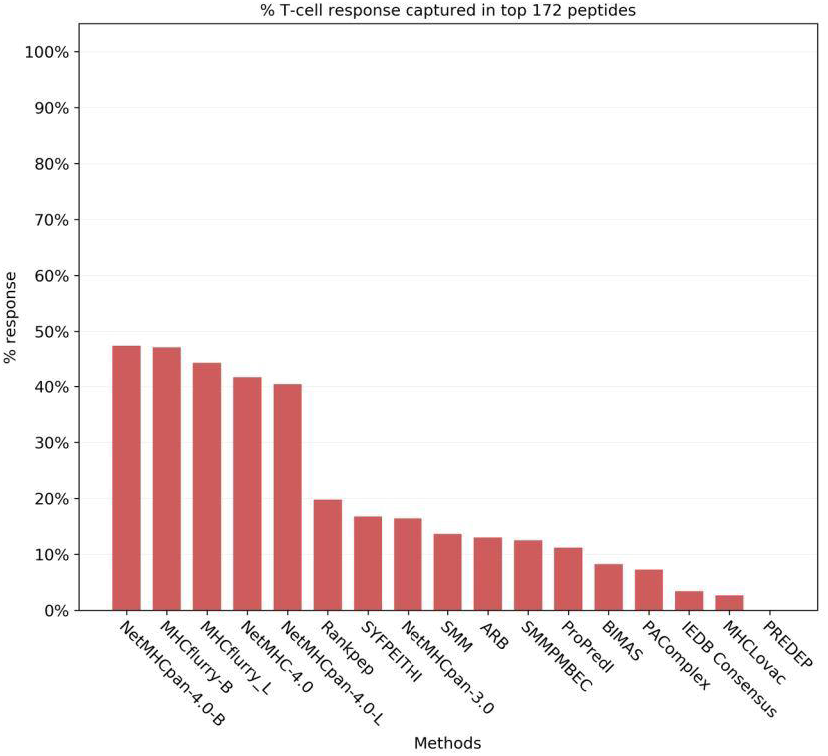
Number of epitopes and the amount of T cell response captured in the top 172 peptides. The number of top peptides was fixed at 172 to match the number of peptides identified by mass-spectrometry. Fig 4A. Number of epitopes captured in the top 172 peptides. Fig 4B. Amount of T cell response captured in the top 172 peptides.

**Table 3.** Number of epitopes and amount of T cell response captured in the top 172 peptides.

| Method | Epitopes captured in<br>top 172 peptides |  | T cell response<br>captured in top 172<br>peptides |  |
| --- | --- | --- | --- | --- |
|  | Count | % | Count | % |
| MHCflurry-L | 36 | 43.37% | 24.92 | 44.36% |
| MHCflurry-B | 36 | 43.37% | 26.51 | 47.18% |
| NetMHCpan-4.0-B | 33 | 40.00% | 26.64 | 47.42% |
| NetMHCpan-4.0-L | 33 | 39.76% | 22.7 | 40.40% |
| NetMHC-4.0 | 31 | 36.86% | 23.5 | 41.83% |
| Rankpep | 22 | 26.51% | 11.1 | 19.75% |
| SYFPEITHI | 16 | 19.73% | 9.43 | 16.78% |
| ProPred1 | 13 | 15.66% | 6.28 | 11.17% |
| ARB | 12 | 14.46% | 7.32 | 13.04% |
| BIMAS | 11 | 13.25% | 4.64 | 8.25% |
| SMM | 11 | 12.67% | 7.65 | 13.61% |
| NetMHCpan-3.0 | 10 | 12.11% | 9.23 | 16.42% |
| SMMPMBEC | 7 | 8.72% | 7.03 | 12.51% |
| PAComplex | 3 | 3.61% | 4.1 | 7.30% |
| IEDB Consensus | 2 | 2.88% | 1.93 | 3.43% |
| MHCLovac | 2 | 2.41% | 1.51 | 2.69% |
| PREDEP | 0 | 0.00% | 0 | 0.00% |

Number of epitopes and amount of T cell response captured in the top 172 peptides. The higher the number of epitopes or amount of T cell response captured, the better the performance of the prediction method. The number of top peptides was fixed at 172 because that was the number of peptides identified by LC-MS/MS.

In addition to the analyses based on the binary classification of peptides (epitopes/non-epitopes), we also evaluated the methods based on the T cell response generated by the peptides, measured as the percentage of IFNγ producing cells in CD8 T cells as a whole (S3 Table). First, we plotted the cumulative fraction of the T cell response accounted for by a given % of the total peptides considered and estimated the overall AUCs for each method with peptides of all lengths and both alleles taken together. Measuring the performance of the prediction methods based on the magnitude of the T cell response covered systematically gave slightly higher performances with overall AUCs ranging from 0.651 to 0.979 (Table 1, Fig 1B). The rankings however were essentially identical, with NetMHCpan-4.0-L again ranking first with an AUC of 0.979 followed by NetMHCpan-4.0-B (0.978) and MHCflurry-L of (0.977). Fig 2E shows the the corresponding curves for 100% peptides and Fig 2F shows the same for 2% peptides. Similar to the analysis we did with epitopes, we also estimated the number of peptides needed to capture 50%, 75% and 90% of the T cell response. The results were essentially same as that of the epitopes at the corresponding percentage levels with some minor exceptions (Table 2, Fig 3D-F). Similarly we also calculated the amount of T cell response captured in the top 172 peptides predicted by each method (Table 3 and Fig 4B). Here NetMHCpan-4.0-B came top with 47.4% of the response and was closely followed by MHCflurry-B with 47.2% T cell response.

### 3.3 Comparing epitope identification by mass-spectrometry and epitope prediction

Next, we wanted to determine how epitope candidates identified experimentally by mass-spectrometry (MS) should be ranked. In the dataset used, a single elution and identification of peptides by LC-MS/MS was done. Rather than treating the outcome of this MS experiment as a binary outcome (ligands being identified or not), we ranked the results based on confidence that the identified hits are accurate, and to test if that enables discriminating hits that turn out to be epitopes from others that do not. We compared the performance of three metrics derived from the MS experiment. First the ProteinPilot confidence score which is obtained from the software used in identification of peptides using MS; second, the number of times a peptide was identified in MS (*i.e.* spectral count); and third, a combined score derived by taking the product of the previous two (S3 Table). When evaluating these three approaches, we found that the number of times the peptide was identified by MS had the best performance with an AUC of 0.674 (AUC of combined score = 0.667, ProteinPilot = 0.503). This shows that the number of times a precursor ion was selected for MS/MS, which is a proxy for the abundance of a peptide, but not the ProteinPilot score, which is an indication of the certainty of the hit, has small but significant predictive power for a peptide to be an actual epitope (p = 0.0001).

Using this score to rank the identified MS ligands, and assigning a score of 0 to all other peptides in the VACV peptide dataset, we could now generate ROC curves in the same way as was done for the prediction approaches, and compare it to the best performing method NetMHCpan-4.0-L. Fig 5A shows the ROC curves for both MS-based and prediction based (NetMHCpan-4.0-L) approaches for 100% FPR and Fig 5B shows the ROC curves up to 2% FPR. The MS based curve had an AUC of 0.898 compared to AUC of 0.977 for NetMHCpan-4.0-L. At the same time, when evaluating how many peptides are needed to be synthesized to capture 50% of the epitopes, the ligand elution data by far outperforms all prediction methods, needing only 0.01% peptides (N = 48), with the best prediction method (NetMHCPan4L) needing 277 peptides. This suggests that, when the intent of a study is to identify all epitopes, and the number of peptides tested is a minor concern, predictions have a better performance, as some fraction of T cell epitopes will be missed in typical ligand-elution experiments. At the same time, when the intent is to identify a small pool of high confidence candidate peptides, MHC ligand elution experiments have a much better performance.

**Fig 5.**
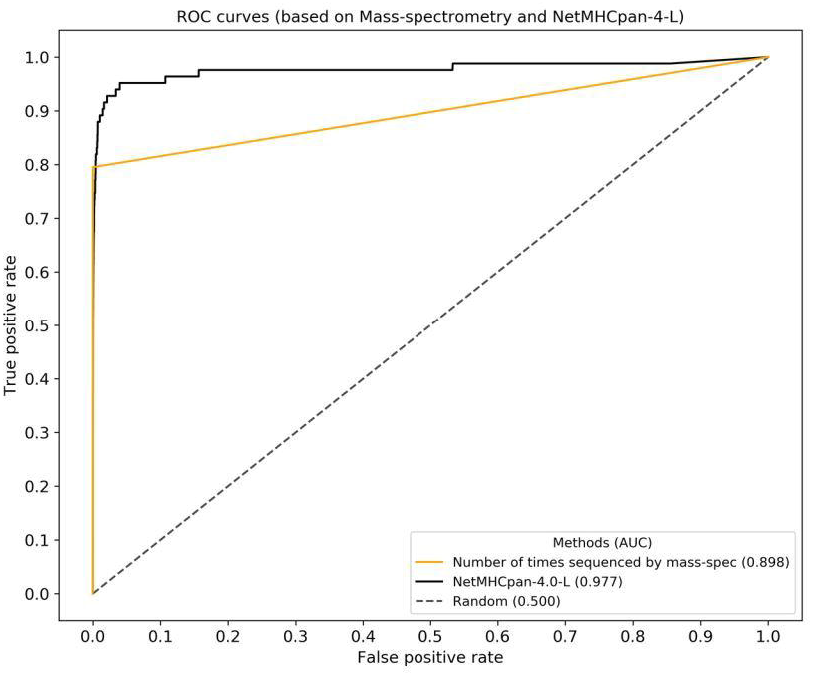
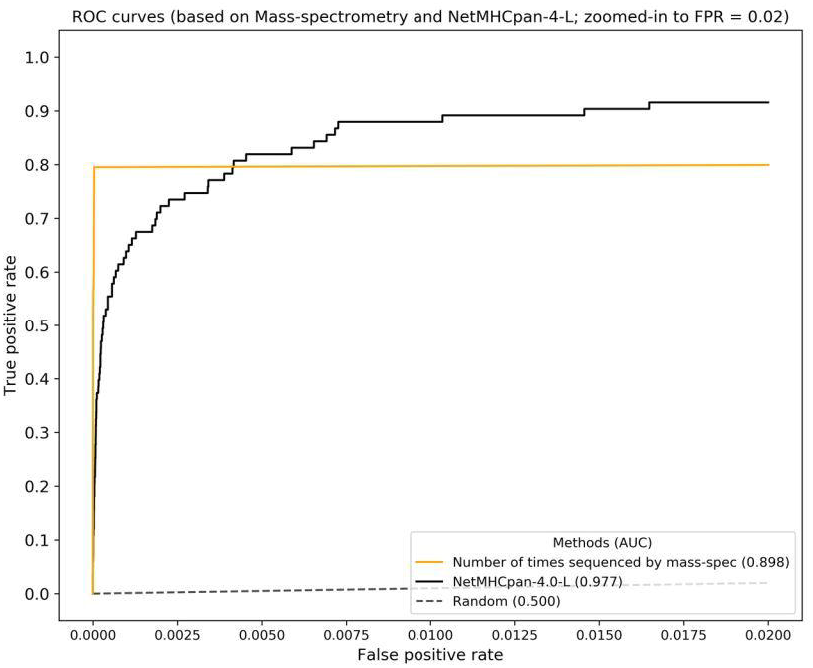
ROC curves comparing epitope candidate selection using mass-spectrometry and prediction approaches. The curves were generated from the number times a precursor ion was selected for MS/MS which acts as a proxy for the abundance of a peptide and represents MS and NetMHCpan-4.0-L prediction scores. Fig 5A. ROC curves comparing epitope candidate selection using mass-spectrometry and prediction approaches. Plot showing 100% FPR. Fig 5B. ROC curves comparing epitope candidate selection using mass-spectrometry and prediction approaches. Plot showing up to 2% FPR.

### 3.4 Comparison of prediction speed

As an independent measure of prediction performance, we wanted to compare the speed with which the different methods could provide their answers. As the initial gathering of predictions involved significant manual troubleshooting, we performed a dedicated speed test, using 5 random amino acid sequences that were 1000 residues long for both H-2D^b^ and H-2K^b^ alleles, and for each method. We used the fastest available online versions of the methods for prediction, for example, RESTful API where available. For some methods, we were unable to quantify prediction times that could be meaningfully compared to the others, and these were excluded from this analysis (for example, MHCflurry server was having memory issues and we could not get the predictions done in a manner consistent with other methods). Out of the 10 methods that we could compare, BIMAS and SYFPEITHI were the fastest with 0.97 and 0.99 seconds per sequence respectively (Fig 6A). On the other end, NetMHCpan-4.0 and NetMHCpan-3.0 were the slowest with average times of 8.53 and 6.30 seconds. We noticed that in general, matrix based methods (BIMAS, SYFPEITHI, RANKPEP, SMM, SMMPMBEC) were significantly faster compared to artificial neural network-based methods (NetMHCpan-4.0, NetMHCpan-3.0, NetMHC-4.0) on average (Fig 6B). The matrix-based methods took an average of 2.07 seconds while the neural network-based methods needed an average of 6.06 seconds per sequence, with the pan-based methods being particularly slow. This indicates a trade-off between prediction performance and speed.

**Fig 6.**
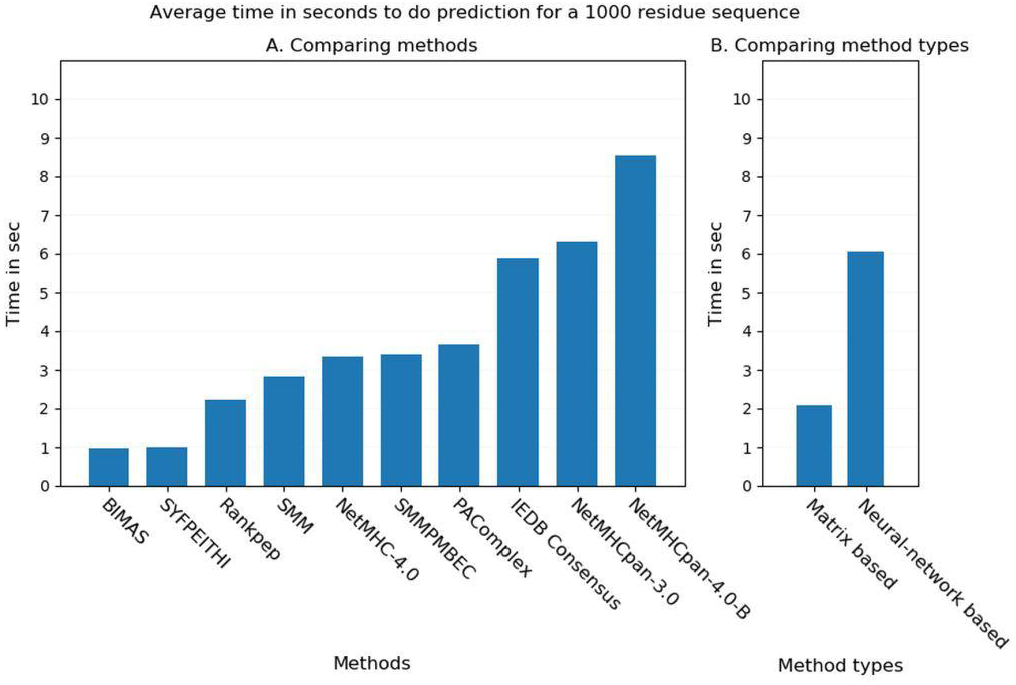
Comparison of prediction speed among the some of the benchmarked methods. The plot shows the average time in seconds taken by the methods for doing epitope prediction for 1000 amino acid residue long sequence. Fig 6A. Comparison of prediction speed among individual methods Fig 6B. Comparison of prediction speed between matrix-based methods and artificial neural network-based methods

## 4. Discussion

In this study we comprehensively evaluated the ability of different prediction methods to identify T cell epitopes. We found that most of the latest methods perform at a very high level, especially the methods developed on artificial neural-network based architectures. In addition, we found that methods that integrated MHC binding and MHC ligand elution data performed better than those trained on MHC binding data alone. And where available, methods that provided two outputs, where one output predicted MHC ligands vs. another that predicted MHC binding, the MHC ligand output score performed better. Based on these results, the IEDB will be updating the default recommended prediction method to NetMHCPan-4.0-L.

Our results highlight the value of integrating both MHC binding and MHC elution data into training prediction algorithms, and confirms that the approach of generating different prediction outputs allows to capture aspects of MHC ligands that is not captured by binding alone, and that these aspects improve T cell epitope predictions [14]. At the same time, the difference in performance is small, highlighting that MHC binding captures nearly all features of peptides that distinguish epitopes from non-epitopes in current prediction methods.

It is also interesting to note that the top 172 peptides captured 40% or more epitopes by the top methods (NetMHCpan-4.0, MHCflurry) (Table 3). This should be viewed against the total amount of peptides in the entire peptidome that could be generated from VACV proteome. It means that the top 0.02% of the peptides could capture 40% of the epitopes and close to 50% of the total immune response (Table 3). Similarly, it took less than 2% of the top peptides predicted by the best methods to capture 90% of the epitopes and T cell response. In the same manner less than 0.04% of peptides captured 50% of the epitopes and T cell response (Table 2). This is relevant because it shows that these methods can significantly reduce the number of peptides needed to be tested in large scale epitope identification studies. Balance between greater coverage (with fewer false negatives) vs. greater specificity (with fewer false positives) that comes with different thresholds and methods has to be made in the context of a specific application. For example, if the goal of a study is to identify patient specific tumor epitopes for a low mutational burden tumor, avoiding false negatives is crucial, as there are few potential targets to begin with. In contrast, if the goal of a study is to identify epitopes that can be used as potential diagnostic markers for a bacterial infection, there will be a plethora of candidates, and avoiding false positives becomes much more important.

A limitation of previous benchmarks is that they either used MHC binding or MHC ligand elution data to evaluate performance, or they use T cell epitope datasets for which it is unclear what constitutes a negative. The dataset we use here is unique in that it comprehensively defines T cell epitopes in a consistent fashion. The downside of this dataset is that it is limited to two murine MHC class I molecules. Future benchmarks on similar datasets for T cell epitopes recognized in humans will be necessary to confirm that the results hold there.

In the process of conducting this benchmark, it became clear that comparing methods that varied in terms of the lengths of peptides they covered introduces difficulties. Developers want to see methods compared on the same datasets, and can refer to the values in Table 1. We strongly advocate that all prediction methods should be evaluated by ranking all possible peptides, which should be extended to ligands from 7 to 15 residues in the case of MHC class I. Method developers should also include guidance on how scores from different length peptides should be compared. That has been done in some cases before [26], but has not been done in others, including in several developed by our own team (SMM, SMMPMBEC).

We want to mention that out of the 172 peptides that were identified by LC-MS/MS, 37 were detected in modified form but were tested for immunogenicity as synthesized unmodified peptides (S3 Table). The caveat is that we do not know to what extent the modification affects binding compared to unmodified form for these peptides or indeed if some modification were artefacts of sample preparation. We therefore repeated the analysis after excluding the peptides identified in modified form and found that the AUCs did not change much and the rankings of the methods remained same except that MHCflurry-B moved ahead of MHCflurry-L (S5 Table).

Although the artificial neural network-based methods were much ahead in performance, they were found to be slower compared to the matrix-based methods. This is expected since artificial neural network-based methods employ more complex algorithms compared to rather linear models used by matrix-based methods. But it should be noted that offline or standalone versions are available for many methods that are significantly faster than the online and API versions. These versions can be run on local computers and users should consider using these standalone versions for doing large scale predictions.

Finally, an important aspect of this benchmark is that we have made all data including prediction results from all benchmarked methods and the code for generating all result metrics and plots publicly available as a pipeline (https://gitlab.com/iedb-tools/cd8-t-cell-epitope-prediction-benchmarking). We believe this will act as a useful resource for streamlined benchmarking process for epitope prediction methods. New prediction method developers can plug in the prediction scores from the new method into this dataset and run the pipeline for side-by-side comparison of their method’s performance with those included in the analysis. The only point to remember is that the developers should exclude this data from the training data for their method. We believe that this benchmark analysis will not only help guide immunologists choose the best epitope prediction methods for their intended use, but will also help method developers evaluate and compare new advances in method development, and provide target metrics to optimize against.

## 5. Author contributions

BP and SP designed the study. SP retrieved predictions, and performed all analysis. NPC, AWP and DCT aided in the interpretation of the MS data in the context of predictions. All authors contributed to the interpretation of the results and writing of the manuscript.

## 7. Supporting information captions

**S1 Table. List of publicly available T cell epitope prediction methods compiled from internet.** There were 44 methods with the executables freely available. This list was further screened for inclusion of the methods in the benchmark analysis based on certain criteria e.g. availability of trained algorithms for the two alleles for which we had data. The last column shows whether the method was included and the reason for exclusion in case it was not included.

**S2 Table. Methods included in this benchmark analysis.** The table shows the methods finally included in the benchmark analysis and their available peptide lengths per allele.

**S3 Table. Peptides tested for T cell response.** The table shows the 220 VACV peptides that were tested for T cell immune response. It includes the 172 peptides that were identified by mass-spectrometry and the additional 48 peptides that were selected from other sources. This table is derived from Croft et al., 2019 (dataset-S1 therein).

**S4 File. The VACV reference proteome used for generating VACV peptides that were used in the analysis.** The proteome was collected from UniProt (Vaccinia virus strain Western Reserve, https://www.uniprot.org/proteomes/UP000000344).

**S5 Table. Overall AUCs after excluding the peptides that were identified in modified form by the LC-MS/MS but tested for T cell response in unmodified form.** The ranking of the methods was same as that with including all peptides with only one exception that MHCflurry-B moved ahead of MHCflurry-L.

## Notes

https://gitlab.com/iedb-tools/cd8-t-cell-epitope-prediction-benchmarking

